# Searching for Sequence Features that Control DNA Cyclizability

**DOI:** 10.1101/2025.01.02.631081

**Authors:** Margarita Gordiychuk, Jonghan Park, Aakash Basu, Taekjip Ha, William Bialek, Yaojun Zhang

## Abstract

The mechanical properties of DNA molecules are crucial in many biological processes, from DNA packaging to transcription regulation. While the mechanics of long DNA typically follow the worm-like chain polymer model, multiple studies have shown that the mechanics of short DNA – at the length scale of DNA-protein interactions – depend strongly on its sequence content. Motivated by recent high-throughput measurements of sequence-dependent DNA cyclizability – the DNA’s tendency to mechanically bend and form a loop, we developed a statistical mechanics approach to systematically explore how cyclizability depends on interactions between individual nucleotides in the sequence. By applying this method to datasets of randomly generated and biologically derived sequences, we identified characteristic sequence features that control DNA cyclizability and extracted the most and least cyclizable sequences, the behavior of which we validated through all-atom molecular dynamics simulations. We found that while highly cyclizable sequences share the same periodic features across datasets, distinct sequence patterns can result in low cyclizability. This work contributes to our understanding of the sequence dependence of DNA mechanics and its role in various biological processes, and has implications for the growing field of DNA nanofabrication.

## I. INTRODUCTION

The double-helical DNA is not just a passive repository of genetic information but an active physical entity that interacts with diverse protein machinery to regulate and control essential biological processes, including genome packaging, transcriptional regulation, DNA repair, and more. Increasingly, the mechanical properties of DNA are recognized to play critical roles in these processes. For example, DNA mechanics can influence where the nucleosomes form and how DNA is packaged within the cell, how DNA interacts with transcription factors and how probable it is to establish enhancer-promoter interactions via DNA looping, as well as how DNA mismatches recruit repair machinery [1–3]. The mechanical behavior of DNA is commonly described by the worm-like chain polymer model, which treats DNA as an elastic rod with a bending persistence length of 50 nm (approximately 150 base pairs) [4–6]. While being successful in capturing the DNA mechanics on long length scales, it predicts that DNA segments shorter than the persistence length are essentially rigid rods with a very low propensity to form loops. However, such predictions were contradicted by observations of prevalent spontaneous large-angle bends in DNA at short length scales (tens of base pairs) [7, 8]. Moreover, single-molecule assays over the years have shown that DNA molecules can exhibit complex mechanical behaviors highly dependent on their nucleotide sequences [9, 10]. Epigenetic modifications (e.g. methylation) and DNA mismatches can further induce non-canonical DNA structures that significantly alter DNA mechanics, especially at short length scales [11].

A key experimental approach to characterizing DNA mechanics is through the measurement of cyclizability, which quantifies a molecule’s ability to bend and form a loop. DNA cyclization has been investigated from a thermodynamic perspective as quantified by the Jacobson–Stockmayer factor [12– Multiple studies have demonstrated that sequence content can significantly affect the J-factor of DNA [9, 15, 16]. However, a systematic characterization of how DNA cyclizability depends on sequence features has been lacking. Recently, a novel high-throughput sequencing-based method called loop-seq was developed, which enabled the simultaneous measurement of cyclizability across hundreds of thousands of sequences [17]. This extensive data set has inspired several deep neural network-based approaches to understand sequence-dependent DNA cyclizability and predict the DNA mechanics at the genome level [18–22]. Here, we develop a statistical mechanics approach to identify sequence features that control DNA cyclizability and predict the most and least cyclizable sequences, which we validate through all-atom molecular dynamics simulations.

## II. RESULTS

### A. High-throughput data on DNA cyclizability

Recent loop-seq experiments [17] have chosen hundreds of thousands of DNA sequences that span yeast genome and random sequences, and estimated the intrinsic cyclizabilities of these sequences by measuring the probability that they close on themselves into a loop, Fig. 1**a**. Briefly, selected variable sequences of length *N* = 50 were flanked by fixed double-stranded adapters (length 25) and single-stranded complementary overhangs (length 10), and immobilized on a bead. The looping reaction was initiated in high-salt buffer for 1 minute after which the unlooped molecules were digested by an exonuclease that only attacks free ends. The remaining population of looped molecules was sequenced and compared with the control ensemble in which the digestion step was omitted. Cyclizability was defined as the log ratio of probabilities for finding the sequences in the looped vs control ensembles. Observations on a small number of sequences showed that this measure correlates very well with direct single-molecule measurements of cyclizability quantified by fluorescence resonance energy transfer (FRET) [17].

**FIG. 1.**
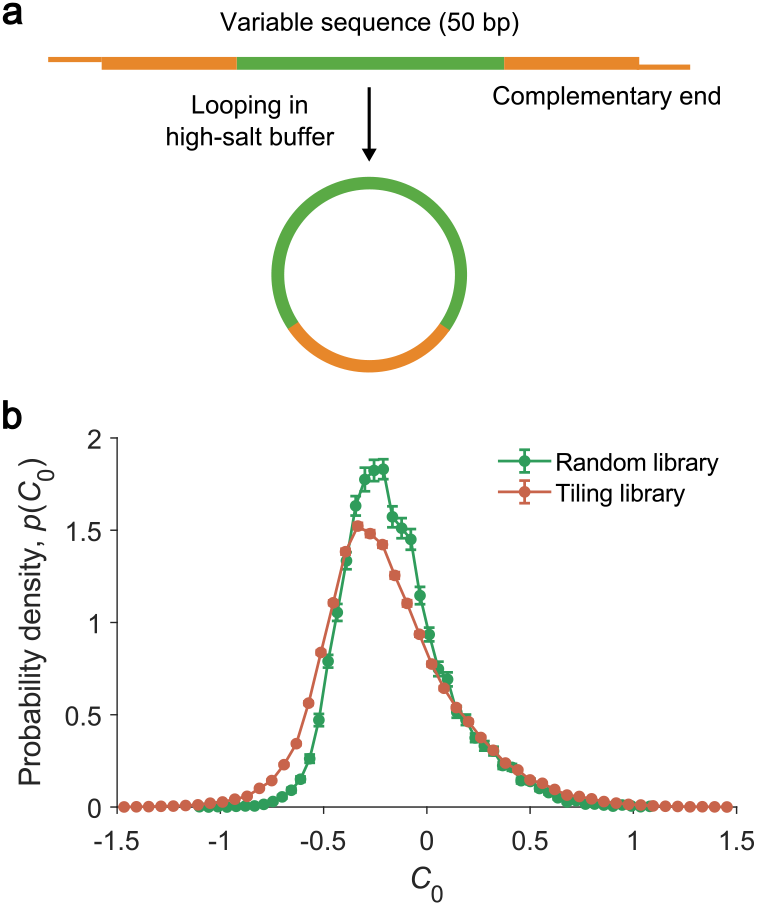
High-throughput measurements of sequence-dependent DNA cyclizability [17]. **a**. Experimental construct. **b**. The probability distribution of the intrinsic cyclizability *C*_0_ for the Random and Tiling libraries. The mean and standard deviation are calculated using random sampling of halves of the data.

The measured cyclizability depends periodically on the location of the bead attachment. The intrinsic cyclizability *C*_0_ was defined as the mean over this variation. The values of *C*_0_ were further refined using a mathematical operation to eliminate the bias introduced by the adapters and overhangs. After these adjustments, *C*_0_ becomes independent of tether location and sequence orientation [22]. The loop-seq data contains multiple libraries of sequences. In this study, we utilize two of them: the Random library, which consists of 12,404 randomly generated sequences, and the Tiling library, which consists of 81,801 sequences tiling around 576 selected genes in the S. cerevisiae genome. The distributions of *C*_0_ across the sequences in the two libraries are shown in Fig. 1**b**.

### B. A statistical mechanics approach to sequence-dependent cyclizability

The intrinsic cyclizability of a sequence is a collective variable determined by contributions from all base sites. To model the cyclizability, we developed a statistical mechanics approach in which cyclizability is systematically expressed in terms of contributions from interactions among an increasing number of bases. To make it concrete, we first convert nucleotide sequences into numbers using one-hot encoding. Specifically, we represent DNA sequences as 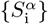, where 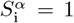 if the base at site i is of type *α*, and 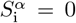 otherwise. The index i = 1, 2, …, *N*, where *N* is the sequence length, and *α* = 1, 2, 3, 4, corresponding to A, T, C, G. In the matrix form, every column of 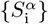 is a unit vector denoting the encoded nucleotide type. For example, 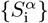 for sequence AGTCGTT…AA is

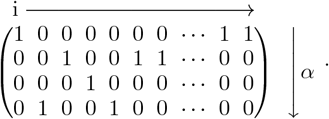

We then express the cyclizability as a cluster expansion, accounting for one-base, two-base, three-base interactions, and so on, in a hierarchical order:

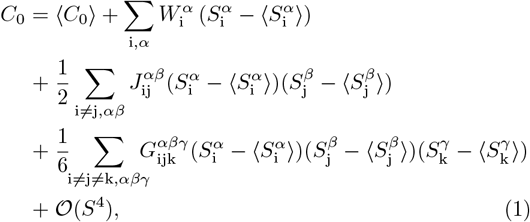

where 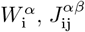, and 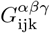 are the coupling constants. In what follows, we explore contributions from these terms order by order, with the goal of building a minimal model that captures the essential sequence features controlling cyclizability.

### C. A linear model

The simplest model for how the cyclizability depends on sequence is linear, where the cyclizability is a weighted sum of the individual nucleotides:

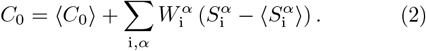

Here, *W* is analogous to the position weight matrices that appear in models of transcription factor binding [23–25]. Without loss of generality, we can set 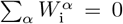 at every site i since 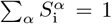 see Methods. If the linear terms are sufficient, then we can extract the elements of *W* by performing a least squares fit of the data with respect to Eq. (2), or, in the particular case when the sequences are random, by computing a correlation function:

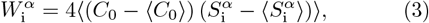

where the average is over all sequences, see Methods for a derivation. Fig. 2 shows the estimate of *W* that results from computing this correlation for the Random library. The results are consistent with *W* = 0, suggesting that there is no linear term in the dependence of *C*_0_ on the sequence. Indeed, when we estimate *W* from 90% of the data and predict *C*_0_ for the remaining 10% using Eq. (2), the correlation coefficient between predictions and measurements is essentially zero, *r* = 0.05 ± 0.03.

**FIG. 2.**
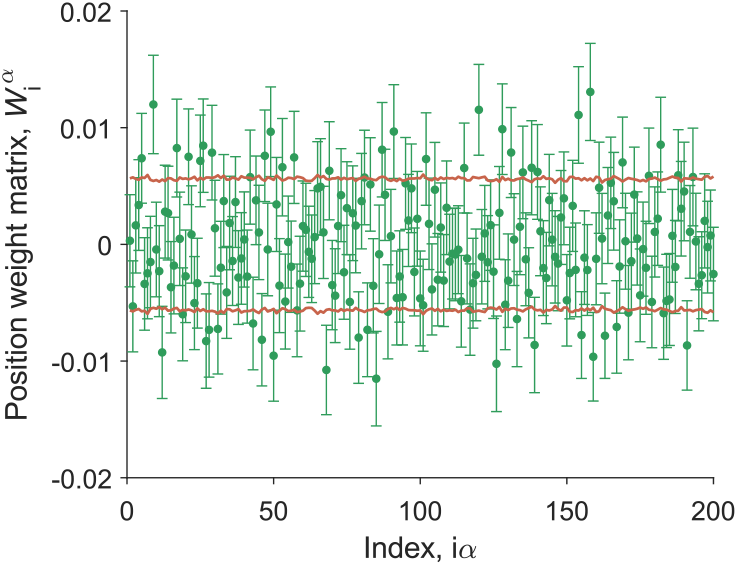
The matrix 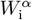 for the Random library computed using Eq. (3). Points: mean and standard deviation across random halves of the data. Lines: ± one standard deviation across random halves of shuffled data. The index i*α* runs from 1 to 200, corresponding to 1A, 1T, 1C, 1G, …, 50A, 50T, 50C, 50G.

### D. A pairwise model

If Eq. (2) doesn’t work, because the data are consistent with *W* = 0, the next simplest model is then a pairwise model, where the intrinsic cyclizability is a weighted sum of interactions between pairs of nucleotides:

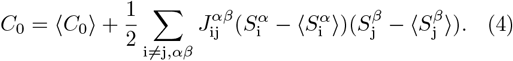

Again, without loss of generality, we can set 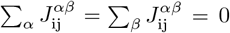 since 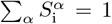, see Methods. As with the linear model, we can extract the elements of *J* by performing a least squares fit of the data with respect to Eq. (4), or, in the case of Random library, directly recover the underlying interaction parameters by computing correlation functions over random sequences:

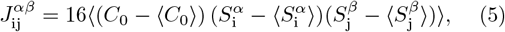

see Methods for a derivation. We can further combine the indices (i, *α*) → i*α* and (j, *β*) → j*β* so that 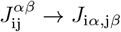. Fig. 3 shows the estimate of *J* from computing the above correlation for the Random library. Compared to the *J* matrix of the shuffled data in Fig. S1, the *J* matrix in Fig. 3 shows clear signals near the diagonal line, highlighting the importance of nearest neighbor interactions. However, when we estimate *J* from 90% of the data and predict *C*_0_ for the remaining 10% using Eq. (4), the correlation coefficient between predictions and measurements is *r* = 0.45 ± 0.02, which is not high.

**FIG. 3.**
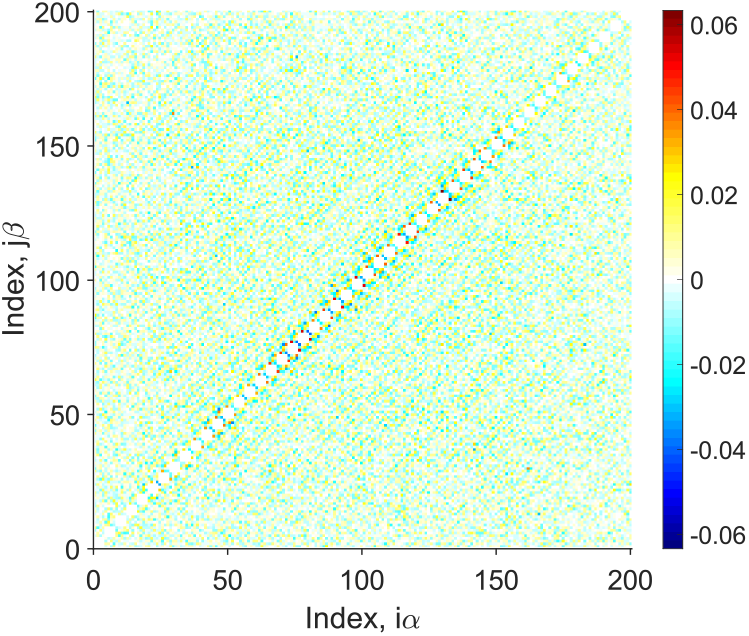
The interaction matrix 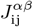 of the pairwise model for the Random library computed from Eq. (5). The indices i*α* and j*β* follow the order 1A, 1T, …, 50C, and 50G.

### E. Physical constraints of the interaction matrix *J*

At first glance, it seems that the pairwise terms are not sufficient to capture the sequence dependence of intrinsic cyclizability, as indicated by the low correlation between the predictions and measurements of the Random Library. Therefore, it seems necessary to include higher-order terms in the model. However, a close inspection reveals that there are 9,800 independent elements in the *J* matrix, comparable to the total number of sequences (12,404) in the Random Library. The limited number of sequences can lead to a low signal-to-noise ratio in the obtained *J* matrix, reducing its predictive power. To improve the predictive performance of the pairwise model, we apply physical constraints to the interaction matrix to reduce the number of variables.

First, 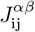 is an interaction parameter between the bases at positions i and j, we expect it to depend on separation but not on absolute position, leading to translational invariance:

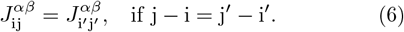

Further, the double helical structure of DNA suggests that a sequence and its reverse complement should have the same cyclizability, leading to reverse complement invariance:

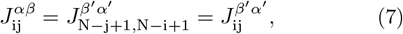

where *α*^*′*^ and *β*^*′*^ denote the complements of *α* and *β* (i.e. A ↔ T and C ↔ G), and translational invariance is used in deriving the last step. We note that by imposing translational and reverse complement invariances, the number of independent elements in the *J* matrix is reduced from 9,800 to 294.

We impose translational invariance by replacing each matrix element of *J* in Fig. 3 with the average of all elements at the same separation of j − i,

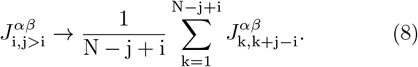

We impose reverse complement invariance by

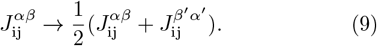

Fig. 4**a** shows the *J* matrix from the Random library after imposing the symmetry conditions. The “symmetrized” *J* not only shows clear signals near the diagonal, suggesting strong nearest-neighbor interactions, but also displays a set of stripes separated at half-helical (∼ 5 bp) and helical (∼ 10 bp) period of DNA, suggesting a role of longer-ranged interactions collectively contribute to DNA cyclizability.

**FIG. 4.**
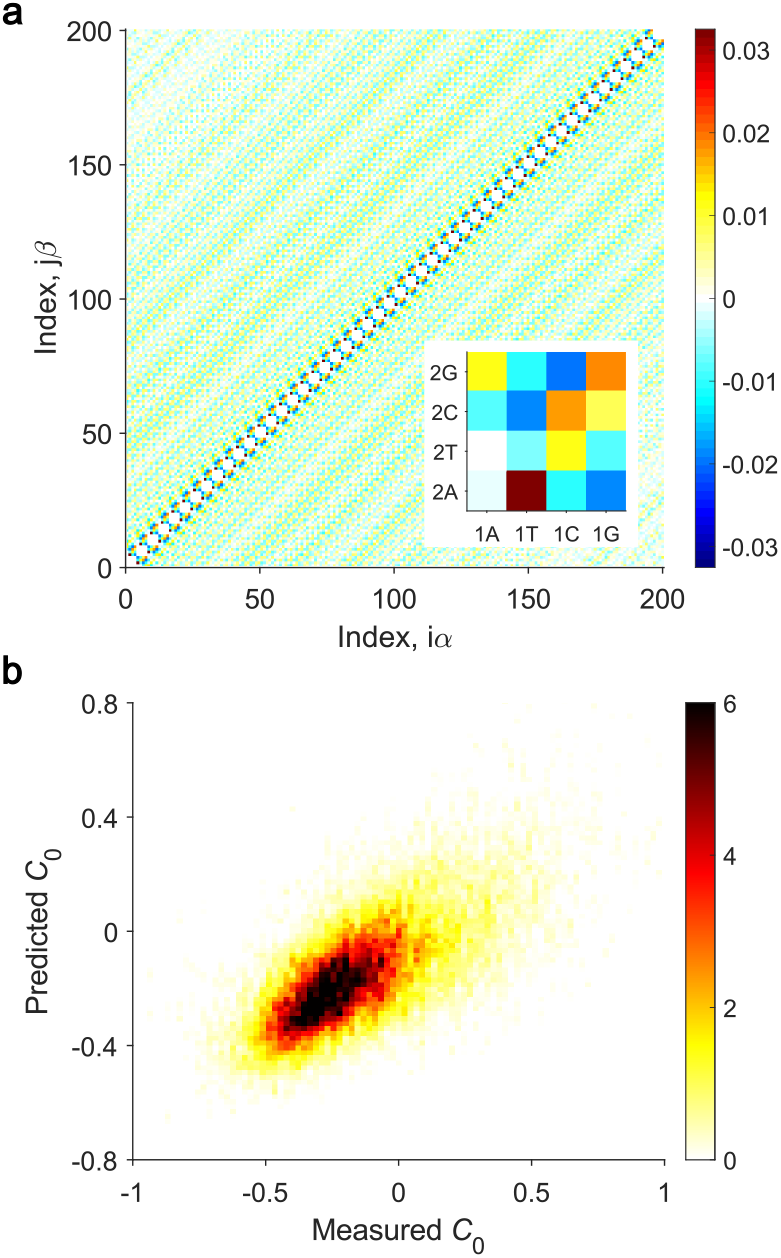
The pairwise model with translational and reverse complement invariances for the sequence-dependence of intrinsic cyclizability. **a**. The interaction matrix 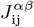 for the Random library after imposing the translational and reverse complement invariances. Inset: nearest-neighbor interaction parameters 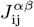 for j − i = 1. **b**. Joint probability distribution of predicted and measured cyclizability *C*_0_ across the ensemble of sequences, Pearson’s correlation coefficient *r* = 0.68 ± 0.02.

Imposing translational and reverse complement invariances raises the signal-to-noise ratio of the inferred *J* matrix, and consequently raises the predictive performance of the model. Predictions vs measurements of intrinsic cyclizability are shown in Fig. 4**b** as a joint density plot, which are obtained from multiple random 90/10 splits of the Random library into training and testing data. The correlation between predictions and measurements is now *r* = 0.68 ± 0.02.

### F. Sequence features that control DNA cyclizability

The predictive performance of the model would likely improve further if we move on to incorporate the three-base interaction terms in Eq. (1). However, the model at the third order involves significantly more parameters than the number of sequences available in the measurements, even after imposing the translational and reverse complement invariances. Therefore, instead of moving to the next order, we search for sequence features that control DNA cyclizability based on the pairwise model in this section.

To identify sequence features, we apply eigenvalue decomposition or principal component analysis to the interaction matrix *J* :

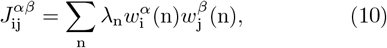

where n is the mode index and the eigenvectors are orthonormal,

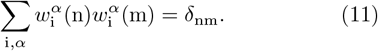

We show in Fig. 5**a** the spectrum of eigenvalues {*λ*_n_} in rank order, and compare with data that have been shuffled to break any correlations between sequence and cyclizability. We first notice that, in both the real and shuffled data, there are some true zero eigenvalues. These arise because we have 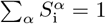 at each site i, by definition. In the shuffled data, we see a spreading of the eigen-values, which arises because the *J* matrix is estimated from a finite sample [26, 27]. Last and most importantly, in the real data, the eigenvalues stand out from the shuffled background with high signal-to-noise ratios at both large positive and negative values, especially at the two largest eigenvalues. We show in Fig. 5**b** the first and last two eigenvectors, which pick out the most and least cyclizable modes of sequence variation. The first two eigenvectors come as a quadrature pair, showing an approximate ten-base periodicity that aligns with the pitch of the double helix DNA. Whereas the last two modes, with their eigenvalues less clearly distinguished from the background noise (Fig. 5**a**), are less clearly periodic and exhibit boundary effects.

**FIG. 5.**
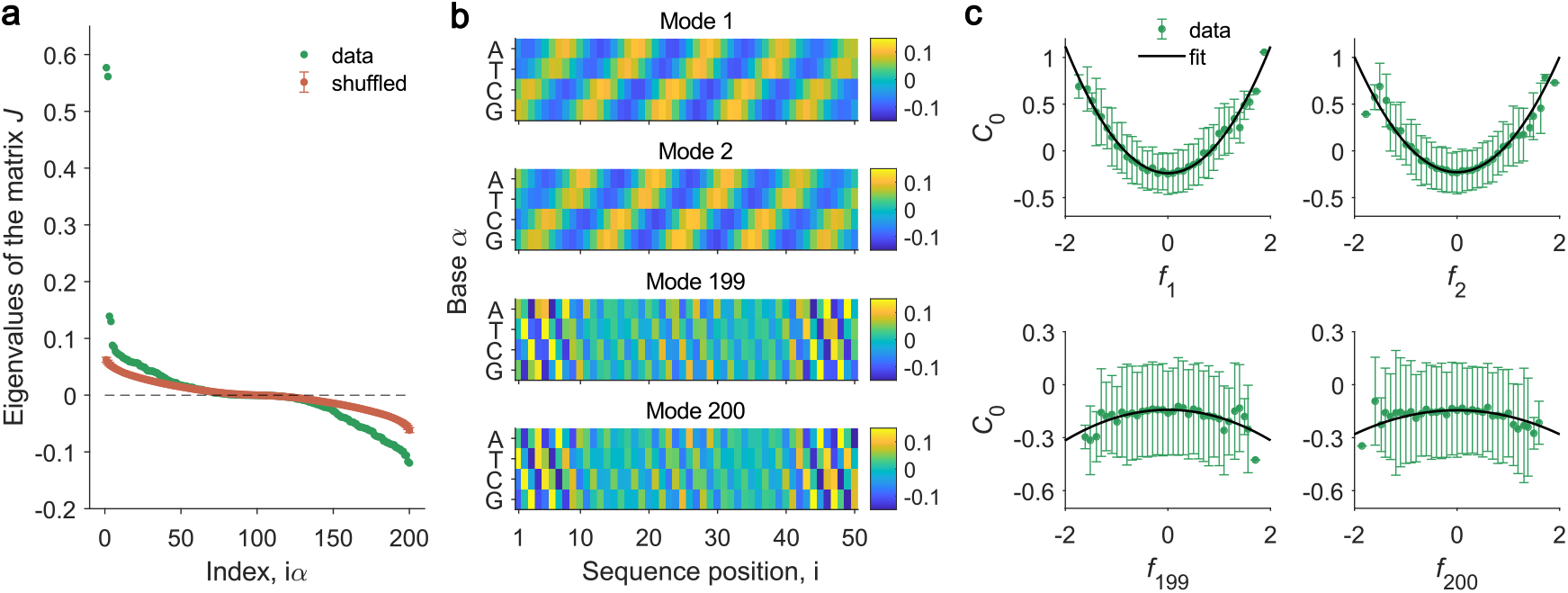
Pairwise interaction model captures sequence features for cyclizability. **a**. Eigenvalues {*λ*_n_} of the interaction matrix *J* from the Random library, ranked in descending order, compared with results from shuffled data of the Random library. Points and error bars in the shuffled data are the means and standard deviations calculated across multiple random 50/50 splittings of the data. **b**. The first two eigenvectors, 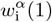 and 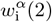, and the last two eigenvectors, 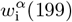 and 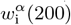, of the matrix *J*. Scale is set by normalization, Eq. (11). **c**. Cyclizability as a function of sequence projection in the feature space. Error bars are standard deviations of the test data set. Lines are quadratic fits as expected from Eq. (14).

We can further decompose the sequence variations into modes defined by the eigenvectors:

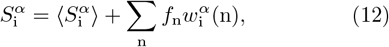

where

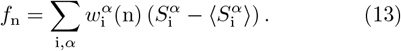

Eq. (4) is then simplified to

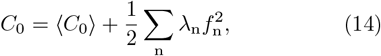

where the sum is over all the modes. In Fig. 5**c**, we show the dependence of the cyclizability *C*_0_ on *f*_n_ at the extremes of the spectrum. To avoid overfitting, we estimate 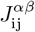 and subsequently the eigenvectors 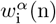 from random half of the sequences, and then probe *C*_0_ vs *f*_n_ in the other half. The mean behavior is quadratic for each mode, consistent with the prediction of Eq. (14).

Equation (14) further enables the separation of contributions from individual modes to cyclizability. Including only the first two modes, i.e. sum over n = 1, 2 in Eq. (14), results in *r* = 0.60 ± 0.02 in the predictive performance, suggesting that these two modes make the largest contribution, but including all modes provides better predictions, Fig. S2.

### G. Comparison of Random and Tiling libraries

Above, we adopted a statistical mechanics approach to analyze cyclizability of sequences in the Random library, which revealed that contributions from linear terms are negligible and that pairwise interactions with imposed invariances are significant for predicting cyclizability. We further derived distinct sequence features for DNA cyclizability from eigenvalue decomposition of the interaction matrix. However, biologically relevant sequences are not random. For example, the yeast genome has a base composition of approximately 31% A, 31% T, 19% C, and 19% G [28], which deviates from the expected 25% of random sequences. To test if our findings from the Random library extend to biological sequences, we analyze a second available dataset, the Tiling library, which comprises sequences from the genome of *S. cerevisiae* [17].

In the analysis of the Random library, we utilized the statistical properties of random sequences to compute the matrices *W* and *J* directly from data, using Eqs. (3) and (5), respectively. These two equations however no longer hold for the Tiling library. Nevertheless, as briefly mentioned before, it is possible to extract the matrices with a least-squares fit applied to Eqs. (2) and (4). To validate the least-squares approach, we first apply it to the Random library. The resulting *W* and *J* matrices match almost exactly those obtained from Eqs. (3) and (5), with Pearson’s correlation coefficients *r* = 0.99 and 0.98, respectively (Fig. S3). We then apply the same approach to the Tiling library. In Fig. S4, we show the extracted *W* which again is consistent with *W* = 0. Correlation coefficient between predictions and measurements of the linear model is *r* = 0.07 ± 0.01, slightly higher than that of the Random library. In Fig. S5, we show the extracted *J* matrix, which shares many similarities with the one from the Random library. Correlation coefficient between predictions and measurements of the pairwise model is *r* = 0.68 ± 0.01, comparable to that of the Random library. For a direct comparison between the Tiling and Random libraries, the matrix elements of *J* for the two libraries were plotted against each other in Fig. 6**a**, which are highly correlated with a correlation coefficient of *r* = 0.90. The most and least cyclizable modes of the two libraries were compared in Fig. 6**b**. While the most cyclizable modes are consistent between the two datasets, the least cyclizable modes are clearly different.

**FIG. 6.**
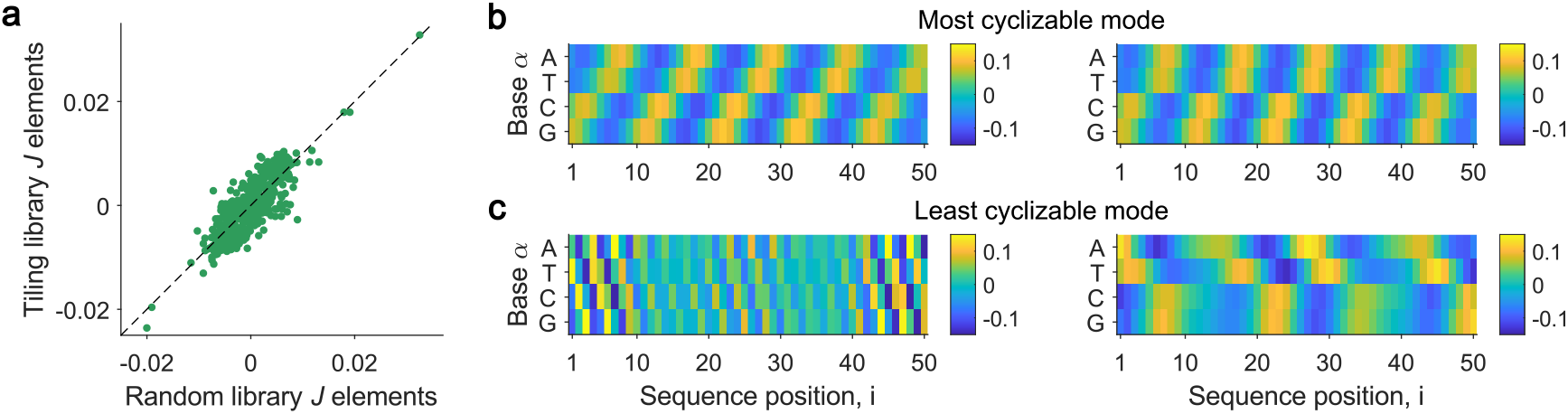
Comparison of Random and Tiling libraries. **a**. Elements of the *J* matrix of the Random library versus the Tiling library, with Pearson’s correlation *r* = 0.90. **b**. The most and least cyclizable modes of the Random library (left) and the Tiling library (right).

We can further construct the most and least cyclizable sequences from the eigenvectors of the two libraries. Because the eigenvectors are orthonormal, increasing the projection of a sequence onto one eigenvector necessarily decreases the projection onto others, the highest and lowest values of *C*_0_ are thus predicted to occur in sequences that have maximal squared projection onto the first and last modes, respectively. Mathematically, this means we look for sequences 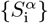 that maximize 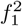 and 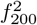 in Eq. (13). We note that because *λ*_1_ and *λ*_2_ (sim-ilarly, *λ*_199_ and *λ*_200_) come almost as a degenerate pair due to translational invariance, sequences that maximize 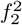 and 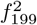 will have comparable cyclizabilities as the most and least cyclizable ones. The predicted most and least cyclizable sequences for the two libraries are listed in Table I. The most cyclizable sequences are identical for both libraries, which consists of 5 − 6 bp tracts of TA-rich segments (e.g. TTAAA) followed by 5 − 6 bp tracts of GC-rich segments (e.g. GGGCCC) periodically. In contrast, the least cyclizable sequence for the Random library consists of short ∼3 bp segments with nucleotides in the order of ATCG, while the sequence for the Tiling library exhibits a periodic structure similar to the most cyclizable sequence but with longer 8 − 10 bp tracts of AT-rich segments where As appear in front of Ts (e.g. AAAAATTTTT).

**TABLE 1.**
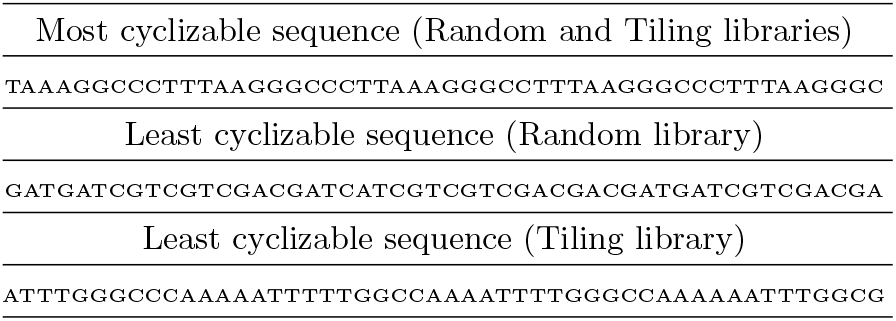
Predicted DNA sequences with highest and lowest intrinsic cyclizabilities.

### H. Molecular dynamics simulations of most and least cyclizable sequences

To further confirm our predictions of the most and least cyclizable sequences in Table I, we performed all-atom molecular dynamics simulations of these sequences in a 1M NaCl solution at constant temperature (300 K) with periodic boundary conditions. The simulations were carried out using the NAMD software [29] with the CHARMM36 force field [30], see Methods for details of the simulation setup. We show in Fig. 7 the snapshots of the DNA at the end of 72-ns simulation runs. The corresponding videos can be found in the Supplementary Material [31]. Remarkably, the most cyclizable sequence curves into a semicircle (Fig. 7**a**), whereas the least cyclizable sequences of both libraries remain straight (Fig. 7**b-c**). Observations from the simulation videos further suggest that the least cyclizable sequence of the Tiling library is slightly more dynamically flexible than that of the Random library. Overall, these results strongly support the predictions of our statistical model.

**FIG. 7.**
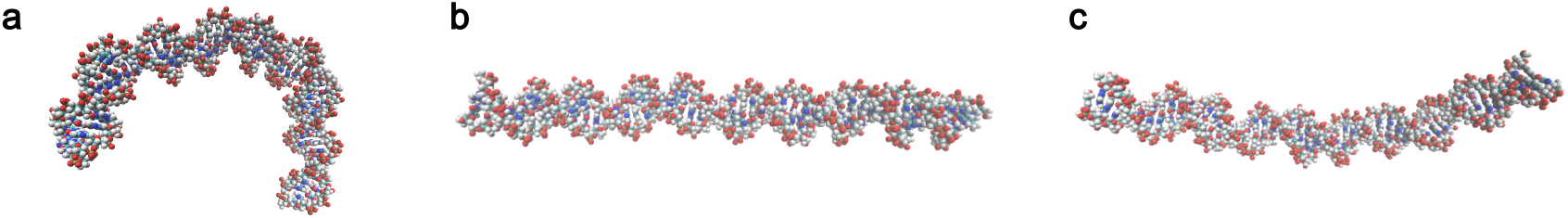
Snapshots of DNA molecules with sequences listed in Table I at the end of 72-ns all-atom simulations performed using NAMD. **a**. Snapshot of the most cyclizable sequence from both the Random and Tiling libraries. **b**. Snapshot of the least cyclizable sequence from the Random library. **c**. Snapshot of the least cyclizable sequence from the Tiling library.

## III. DISCUSSION

The mechanical properties of DNA play an important role in many biological processes. Recent experiments have revealed that the mechanics of short DNA molecules depends strongly on their sequences. However, a systematic characterization of the sequence features that control DNA mechanics has been lacking. In this work, we developed a statistical mechanics approach to analyze high-throughput measurements of DNA cyclizability of randomly generated and biologically derived sequences. We found that a minimal model with pairwise interactions between nucleotides is sufficient to account for the sequence dependence of cyclizability, leading to the prediction of the most and least cyclizable sequences. We finally validated these findings through all-atom molecular dynamics simulations of predicted sequences.

How do our findings compare to those in the literature? Early studies on the sequence dependence of DNA mechanics primarily focused on the influence of dinucleotide pairs due to limited data [32, 33]. Recent development in high-throughput methods have enabled investigations that extend nucleotide interactions to longer length scales. In many respects, our findings align with the consensus knowledge in the field [2, 34]. To be specific, our predicted most cyclizable sequence is consistent with findings from recent SELEX experiments [35], where the selected highly loopable sequences are enriched with alternating short segments of A/T and C/G sepa-rated by half a helical pitch of DNA. Mechanistically, both dynamic flexibility and static bending contribute to the cyclizability of a sequence. Our all-atom simulations suggest that the increased cyclizability of the most cyclizable sequence is mainly due to constructive bending toward one direction, driven by intrinsic curvatures encoded in the sequence. This is analogous to the previously reported globally curved structures formed by sequences with in-phased A-tract repeats [36– Our predicted least cyclizable sequence from the Random library is enriched in CpG steps, which was reported to increase the rigidity of the sequence [39, 40]. The least cyclizable sequence from the Tiling library is enriched in out-phased A-tracts, also known to be very rigid [9].

One limitation of this work is that the predictive power of the pairwise model is lower compared to recent deep neural network models. The Pearson’s correlation coefficient between prediction and measurement is *r* ∼ 0.7 for the pairwise model, whereas *r* ∼ 0.8 − 0.9 for deep neural networks [18–21] and notably *r* = 0.96 in the most recent one [22]. Consequently, the sequence features extracted from the pairwise model may exhibit a similar level of inaccuracy. While we expect that incorporating three-body interactions will enhance the model’s predictive performance, our ability to extract robust third-order parameters is limited by the number of sequences in the datasets. One potential solution is to leverage the high predictive power of deep neural networks to produce larger “synthetic” datasets. Our method can then be employed to crack the “black box” and extract relevant sequence fea-tures.

Moving beyond sequence-dependent DNA mechanics, our method establishes a general framework to address a broad class of problems, where the input lives in a high-dimensional space and the output represents a functional property of the system. With the continued advancement of high-throughput technologies, we anticipate the method developed here to have broad applications. Representative problems in this class include the search for relevant sequence features of DNA, RNA, and proteins that determine their structures, interactions, and functions, as well as the search for relevant stimulus features that govern the responses of sensory neurons.

## IV. METHODS

### A. Details of theoretical methods

1. Justification of setting 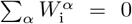. Assume we find the matrix 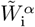 for the linear model, however 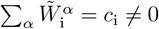. Let 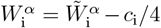, the second term on the right of Eq. (2) becomes:

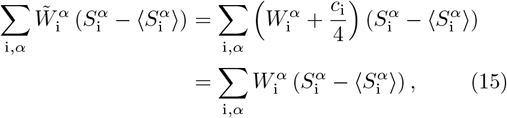

where the *c*_i_ term is gone because 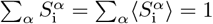 for any given i. Therefore, 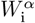 is also a solution to the linear model and we have 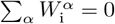.
2. Justification of setting 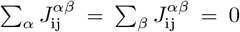. Similar to the linear model, assume we find the matrix 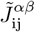 for the pairwise model, however 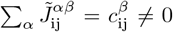 and 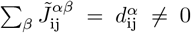. Let 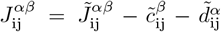, where 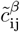 and 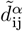 are the solutions of 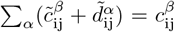 and 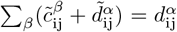, the second term on the right of Eq. (4) becomes:

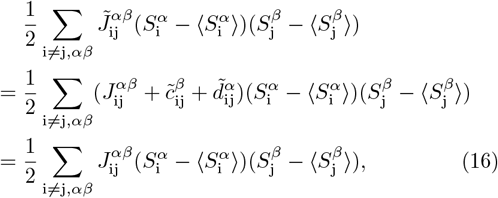

where again the 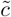 and 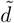 terms are gone because 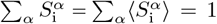. Therefore, 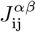 is also a solution to the pairwise model and we have 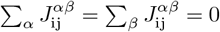.
3. Derivation of Eq. (3) for the Random library. The sequences in the Random library were chosen randomly from a uniform distribution, therefore, we have 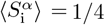 and

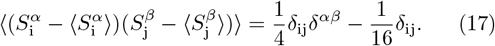 The correlation function between *C*_0_ in Eq. (2) and 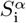 is then

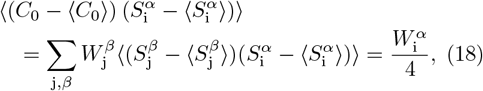

which is Eq. (3).
4. Derivation of Eq. (5) for the Random library. Similar to the above derivation, the correlation function be-tween *C*_0_ in Eq. (4), 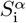, and 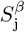 are

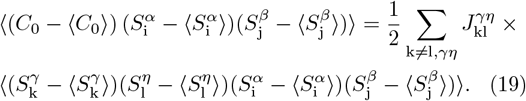 The second line in the above equation is nonzero only if k = i and l = j (case I) or k = j and l = i (case II). For case I, we have

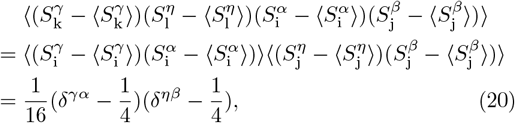

where Eq. (17) was used to derive the last line. Similarly, for case II, we have

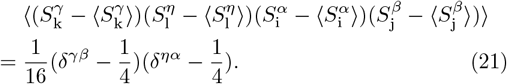 Substituting results in Eqs. (20) and (21) into Eq. (19), we have

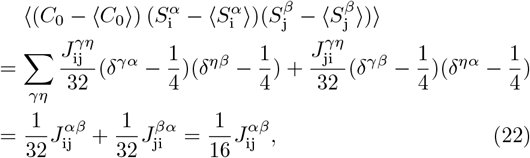

which is Eq. (5).

### B. Details of molecular dynamics simulations

All-atom molecular dynamics simulations of 50-bp DNA sequences in Table I were conducted following the outlined steps below, which were adapted from a previously developed pipeline for quantifying the flexibility of DNA molecules [11].

1. Construction of initial simulation files. The initial structure of the dsDNA molecule was built from the nucleotide sequence using the web 3DNA 2.0 server [41], which generated the starting structure in the B-DNA form with the double helix arranged in a straight line. The resulting PDB file was loaded into the VMD program [42], where the DNA molecule was solvated and ionized. To ensure that the water box dimensions were sufficiently large to accommodate the DNA, we used a 220 Å × 80 Å × 80 Å solvation box for the 50-bp DNA (contour length 170 Å). We next added 1 M concentration of NaCl to neutralize the system and mimic the experimental salt conditions. To prevent the DNA ends from fraying during simulation, we introduced two additional bonds between the ends of the two DNA strands. The PSF and PDB files containing the topological and structural information of all molecules inside the simulation box were then exported from VMD.
2. Energy minimization and equilibration. The solvated and ionized DNA molecule was simulated in NAMD [29] with a 2-fs integration time step and periodic boundary conditions. During the initial energy minimization and equilibration steps, we restrained the DNA molecule to its original coordinates, which allows the surrounding water and ions to equilibrate without disrupting the double-stranded DNA by large forces involved in these processes. Energy minimization was conducted for 10,000 fs to resolve steric clashes and bad contacts. This was followed by a 1-ns constant-volume simulation and a 10-ns constant-pressure (1 bar) simulation both at 298 K to ensure the proper equilibration of the solvent and ions.
3. Final simulation run and trajectory recording. After energy minimization and equilibration, we conducted the final constant-pressure simulation with unrestrained DNA for around 72 ns. Trajectories of all molecules were saved every 9.6 ps for a total of about 7400 recordings. Snapshots and videos of the DNA molecules were built in VMD using these trajectories. The snapshots of DNA at the end of simulations were shown in Fig. 7.

## ACKNOWLEDGMENTS

MG and YZ were supported by a startup fund at Johns Hopkins University. WB and YZ were supported in part by the National Science Foundation through the Center for the Physics of Biological Function (PHY–1734030) and Grant PHY–1607612. AB was a Simons Foundation Fellow of the Life Sciences Research Foundation, and TH is an Investigator with the Howard Hughes Medical Institute.

