## Supplemental Material for "Searching for Sequence Features that Control DNA Cyclizability"

### SUPPLEMENTARY FIGURES

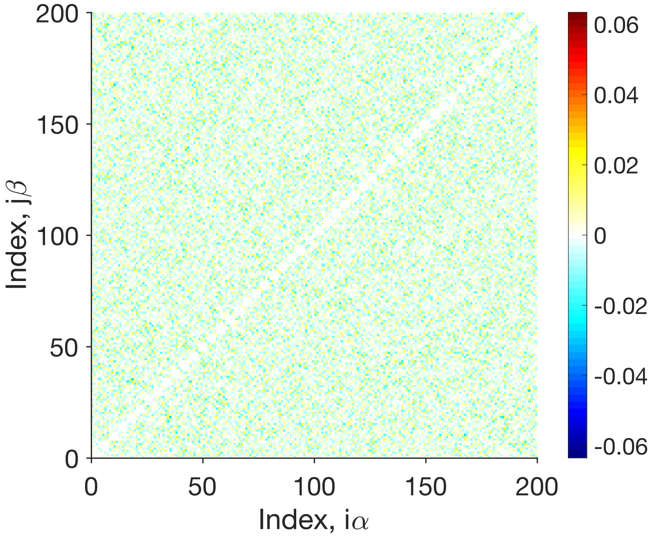


FIG. S1. The interaction matrix $J$ of the pairwise model for the shuffled data from the Random library.

*

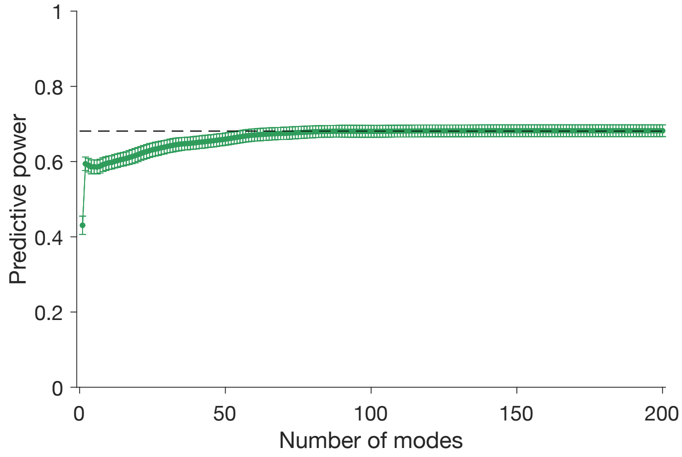


FIG. S2. The predictive performance of the pairwise model as a function of the number of modes included for the Random library. The modes are ranked in descending order based on the absolute values of their eigenvalues.

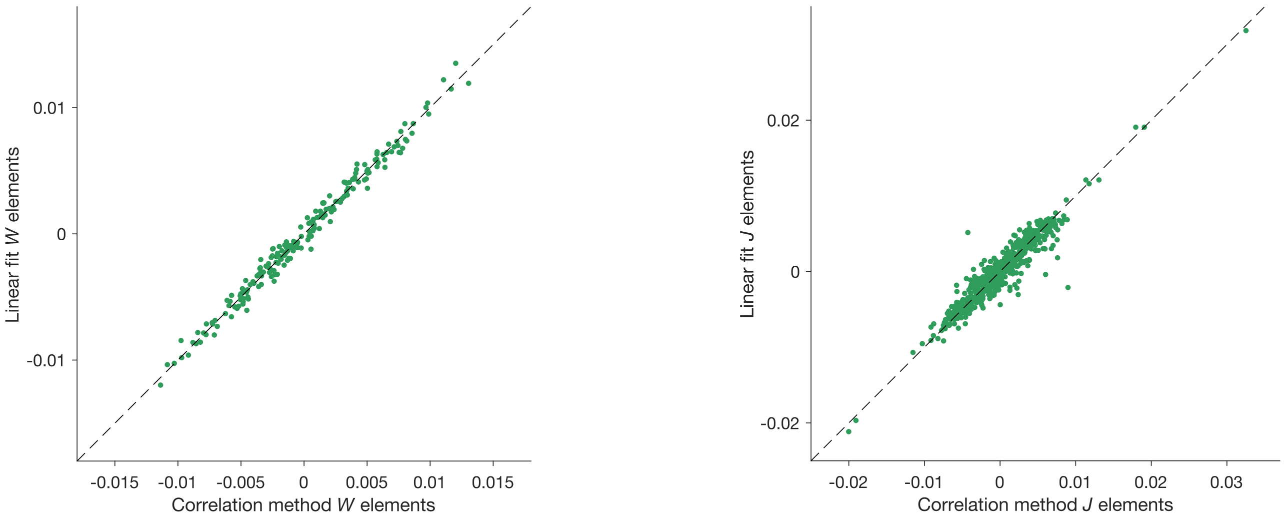


FIG. S3. Comparison of matrix elements obtained using the correlation function approach and the least-squares approach for the Random library. Left: matrix $W$ of the linear model; Right: matrix $J$ of the pairwise model.

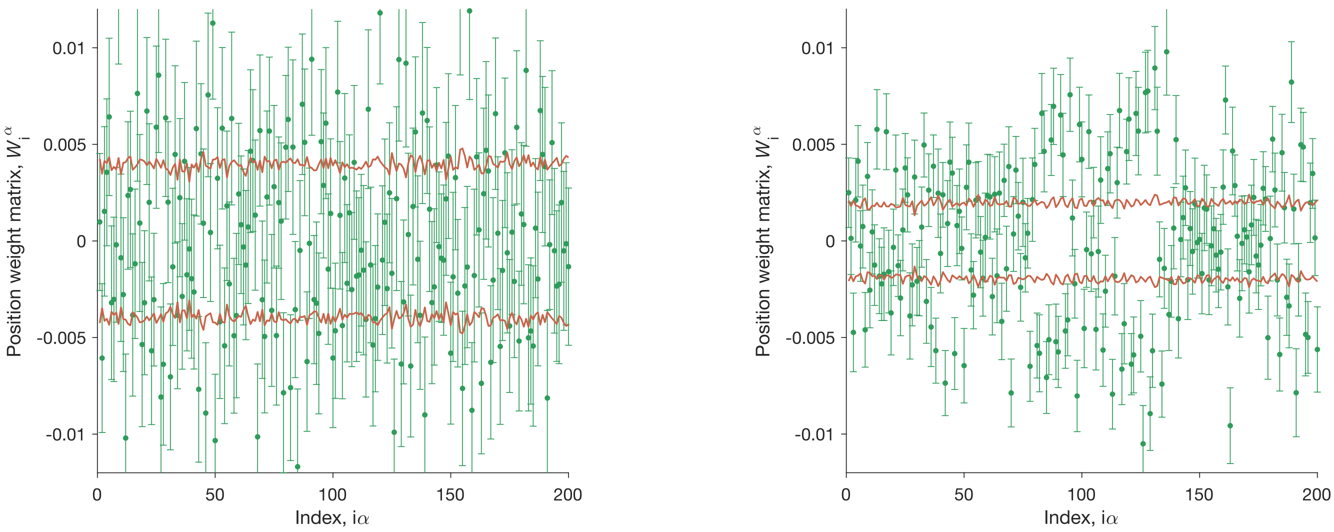


FIG. S4. The matrix $W$ of the linear model extracted from a least-squares fit for the Random Library (left) and the Tiling Library (right). Points: mean and standard deviation across random halves of the data. Lines: ± one standard deviation across random halves of shuffled data.

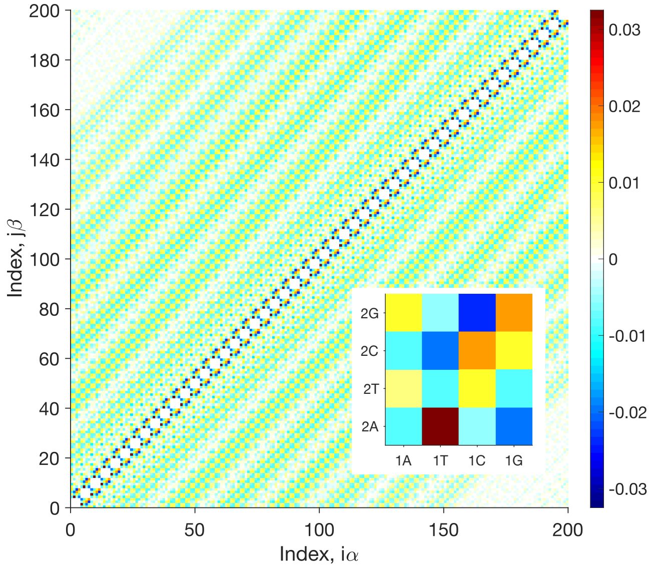


FIG. S5. The interaction matrix $J$ of the pairwise model for the Tiling Library extracted from a least-squares fit. Inset: nearest-neighbor interaction parameters.
